# Prior flavivirus immunity skews the yellow fever vaccine response to expand cross-reactive antibodies with increased risk of antibody dependent enhancement of Zika and dengue virus infection

**DOI:** 10.1101/2023.05.07.539594

**Authors:** Antonio Santos-Peral, Fabian Luppa, Sebastian Goresch, Elena Nikolova, Magdalena Zaucha, Lisa Lehmann, Frank Dahlstroem, Hadi Karimzadeh, Beate M Kummerer, Julia Thorn-Seshold, Elena Winheim, Gerhard Dobler, Michael Hoelscher, Stefan Endres, Anne B Krug, Michael Pritsch, Giovanna Barba-Spaeth, Simon Rothenfusser

**Author notes:** Contributed equally (Giovanna Barba-Spaeth, Simon Rothenfusser).

## Abstract

Human pathogenic flaviviruses pose a significant health concern and vaccination is the most effective instrument to control their circulation. How pre-existing immunity to antigenically related viruses modulates immunization outcome remains poorly understood. In this study, we evaluated the effect of vaccination against tick-borne encephalitis virus (TBEV) on the epitope immunodominance and immunogenicity of the yellow fever 17D vaccine (YF17D) in a cohort of 250 human vaccinees.

Following YF17D vaccination, all study participants seroconverted and generated protective neutralizing antibody titers. At day 28, TBEV pre-immunity did not affect the polyclonal neutralizing response which largely depended on the IgM fraction. We found that sera from TBEV-immunized individuals enhanced YF17D vaccine virus infection via antibody-dependent enhancement (ADE). Upon vaccination, individuals with TBEV pre-immunity had higher concentrations of cross-reactive IgG antibodies with limited neutralizing capacity against YF17D whereas vaccinees without prior flavivirus exposure showed a non-cross-reacting response. Using a set of recombinant YF17D envelope protein mutants displaying different epitopes, we identified quaternary epitopes as the primary target of neutralizing antibodies. Sequential immunizations redirected the IgG response towards the pan-flavivirus fusion loop epitope (FLE) with the potential to mediate enhancement of dengue and Zika virus infections whereas TBEV naïve individuals elicited an IgG response directed towards neutralizing epitopes without an enhancing effect.

We propose that the YF17D vaccine effectively conceals the FLE and primes a neutralizing IgG response in individuals with no prior flavivirus exposure. In contrast, the response in TBEV-experienced recipients favors weakly-neutralizing, cross-reactive epitopes potentially increasing the risk of severe dengue and Zika disease due to ADE.

## Introduction

Human pathogenic flaviviruses comprise over thirty antigenically related viruses (1). Tick-borne encephalitis virus (TBEV), mosquito-borne yellow fever (YFV), dengue (DENV), Zika (ZIKV), West Nile (WNV) or Japanese encephalitis virus (JEV) are flaviviruses with the potential to cause severe disease, representing a leading cause of morbidity and mortality worldwide infecting up to 400 million people annually (2). Flaviviruses are distributed globally, their arthropod vectors are found on all continents and they are rapidly spreading due to international trade and travel, poorly planned urbanization and ecological and climate changes (3,4). Their global distribution, high prevalence and increasing vaccination coverage result in a rising number of individuals with immune experience to flaviviruses. Hence, cross-reactive immunity at the time of vaccination or natural infection with another member of the *Flaviviridae* family is likely to occur.

Flaviviruses share a similar structure, mode of cell entry and mechanisms of maturation and assembly. They are spherical enveloped viruses of about 50 nm in diameter containing a single-stranded, positive-sense RNA genome of about 11,000 nucleotides encoding for a polyprotein that is post-translationally cleaved into three structural proteins: capsid, pre-membrane (prM) and envelope (E), and seven non-structural proteins. In mature virions, the structural proteins are inserted into the host-derived lipid bilayer in an icosahedral architecture where 180 units of the E protein, organized in 90 head-to-tail homodimers, cover the surface of the virion. The E protein is the main target of the antibody response, therefore its structure and dynamics define the epitope landscape of the virus (5,6). The E protein consists of three structurally defined domains (DI, DII and DIII), connected to two transmembrane domains by three stem helices. DII contains the highly conserved hydrophobic fusion loop (FL) (7). A significant portion of the antibody response is directed towards the fusion loop epitope (FLE), which is generally concealed in the dimeric arrangement of the E protein. Although FLE is immunodominant and cross-reactive, its poor accessibility renders fusion loop antibodies (FL-ab) weakly neutralizing (7–9). For effective anti-flavivirus immunity, a relevant fraction of the humoral response targets the E dimer epitope (EDE). This quaternary epitope encompasses regions in DI and DIII of one protomer and DII of the opposing protomer requiring a dimer subunit to be exposed. Upon binding, EDE antibodies crosslink the dimer preventing the conformational changes necessary for fusion. EDE antibodies have been precisely mapped for dengue and Zika viruses (10–12).

Notably, antibody cross-reactivity has been described even among distantly related flaviviruses, which can impact the immune response and clinical course in secondary infections (13,14). Indeed, cross-reactive antibodies can facilitate virus entry via Fcγ-receptor-mediated phagocytosis in a process known as antibody-dependent enhancement (ADE) (15). ADE is associated with severe disease in secondary DENV infections in humans (16–19). Likewise, pre-acquired cross-reactive immunity can impact vaccine responses in different ways. If it propitiates virus neutralization, faster clearance or epitope masking it might lead to suboptimal boosting of immunity (20). Alternatively, cross-reactive antibodies can enhance productive infection of antigen presenting cells via ADE leading to increased immune response (21,22). Lastly, immune imprinting from an earlier immunization may hamper mounting an adequate response against a new antigenic challenge (23).

The YF17D vaccine induces life-long protective immunity and is considered one of the most effective vaccines ever developed (24–27). Prior studies highlighted dimer-dependent epitopes and the FL-proximal region of the E protein as important neutralizing sites for YFV (28–31). In sera of human vaccinees, the response predominantly targets DII with only a minor fraction targeting DIII (32,33). Inter-or intra-dimer epitopes have been structurally mapped for JEV, ZIKV, DENV, and WNV but not for YFV (7) and studies in humans have been restricted to E monomer moieties which account for only a small part of the neutralizing response (33). YF17D is often administered to individuals with flavivirus immune experience. Although the effects of sequential exposures to different flavivirus infections or vaccinations on the immune response have been previously investigated (21,34–37) YF17D-induced antibody specificities and vaccine immunogenicity in the context of pre-existing immunity remain insufficiently characterized.

In this study, we investigated in a longitudinal cohort of 250 participants how pre-existing immunity to TBEV, an antigenically related but phylogenetically distant flavivirus, affects the immunogenicity and epitope immunodominance of the YF17D vaccine. We found that in flavivirus-naïve individuals YF17D elicits non-cross-reactive but efficiently neutralizing antibodies. In contrast, the antibody response in TBEV-experienced individuals is skewed towards the cross-reactive FLE. We further demonstrate *in vitro* that this response results in ADE of dengue and Zika infection.

## Results

### Results 1. YF17D vaccination induces similar neutralizing antibody titers but boosts a poor-neutralizing IgG response in TBEV-experienced individuals

To investigate the effects of pre-existing cross-reactive immunity on the response to the live YF17D vaccine, we examined a longitudinal cohort of 250 YF17D healthy young vaccinees grouped based on their previous immunization with the inactivated TBEV vaccine. Given that TBEV vaccination is recommended in the region where this study was conducted, a representative fraction of participants self-reported a history of TBEV vaccination prior (>4 weeks) to study inclusion (n=162; 64.8%). TBEV pre-immunity was verified through positive results for TBEV neutralization by plaque reduction neutralization assay (PRNT) and for anti-TBEV IgG using enzyme-linked immunosorbent assay (ELISA). Individuals with discrepancies between self-reported vaccination status and serological assays were excluded from analysis. The final study cohort comprises 139 participants pre-vaccinated against TBEV and 56 TBEV-naïve individuals (Fig. 1A-C). The exact strain, number of doses, and timing of TBEV vaccinations had not been documented. Consequently, TBEV-experienced donors showed heterogeneity in their IgG and neutralizing titers.

**Figure 1.**
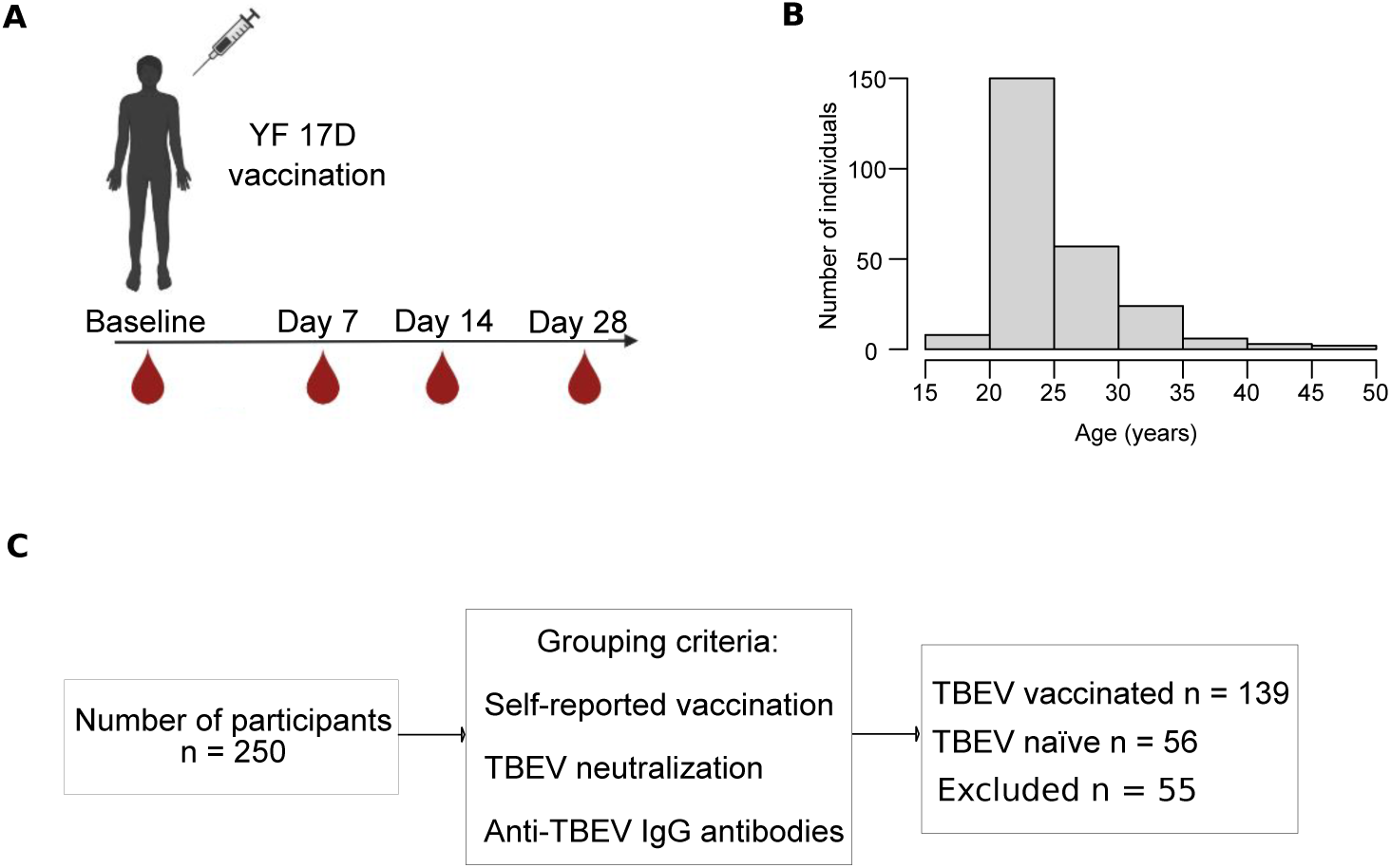
Study subjects overview. A) Diagram representing the longitudinal PBMC, serum and plasma sample collection of 250 participants. B) Histogram depicting age ranges of the study participants. C) Flow chart of cohort members grouping according to TBEV pre-vaccination status. 139 individuals self-reported having received at least one TBEV-vaccine dose and contained neutralizing antibodies and anti-TBEV IgG at baseline. 56 individuals were naïve to flavivirus immunity and were classified based on a self-reported negative vaccination history validated by the absence of detectable TBEV IgG and neutralizing capacity at baseline. 55 individuals were excluded for discrepancies between self-reported vaccination status and serological assays.

The neutralizing antibody titer against YF17D, commonly used as correlate of vaccine-induced protection, was equally strong at day 28 post-vaccination (pv) in flavivirus naïve and experienced individuals. This result indicates that TBEV pre-immunity does not impair the neutralizing antibody response to YF17D (Fig. 2A). Conversely, YF17D immunization did not alter the neutralizing activity against TBEV indicating that the YF17D vaccine does not generate cross-neutralizing antibodies to TBEV (Fig. 2B).

**Figure 2.**
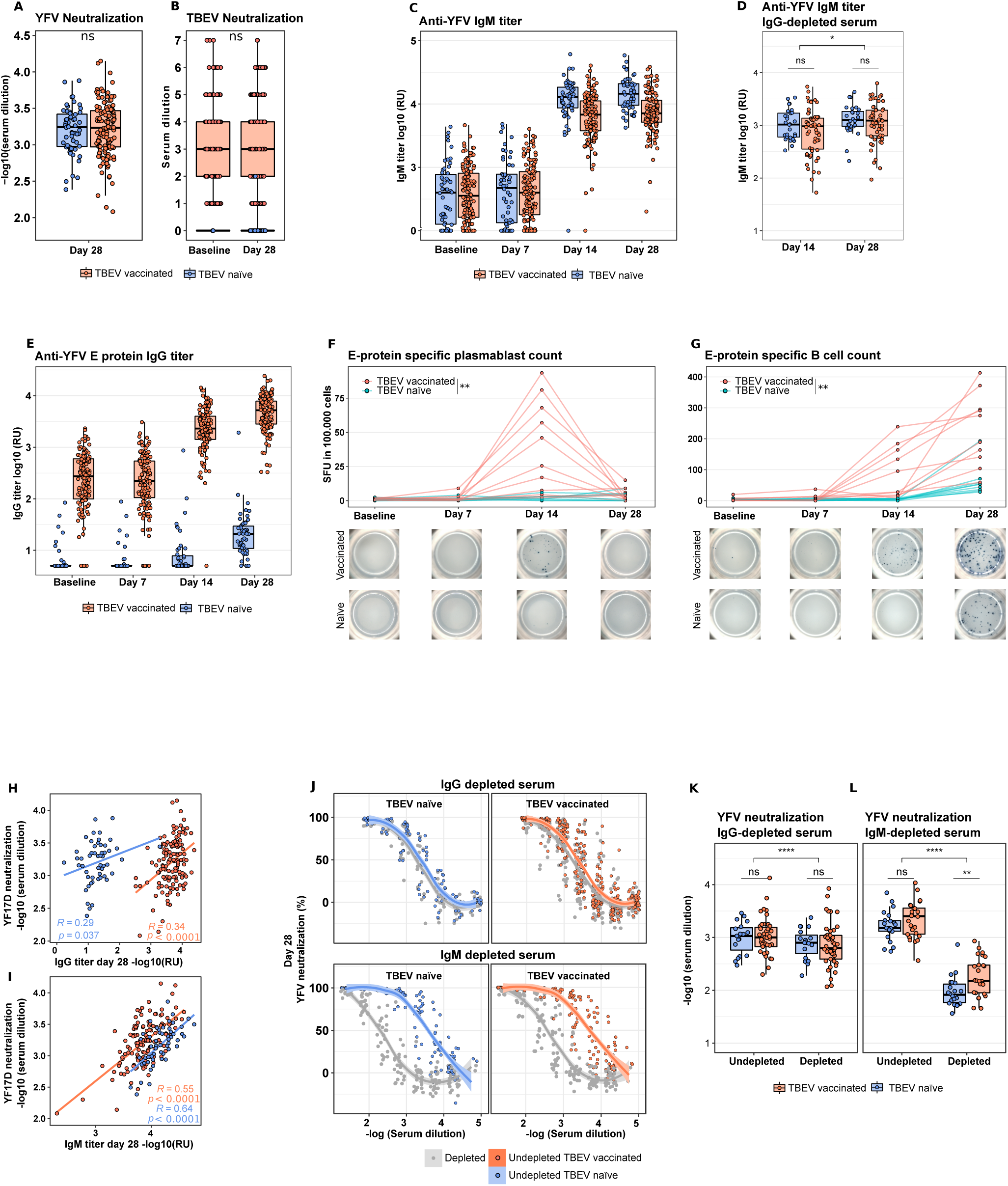
Comparison of the YF17D-induced antibody and B cell responses between TBEV vaccinated and naïve participants. A) YF17D 80 % neutralization titer at day 28 post vaccination. B) TBEV 80 % neutralization titer before and 28 days after YF17D vaccination. C) Longitudinal YF17D virion-specific IgM response in donors’ sera. D) YF17D virion-specific IgM titer at day 14 and 28 in IgG depleted serum. E) Longitudinal YF17D anti-E protein-specific IgG titers. F-G) Plasmablasts and total sE-specific longitudinal B-cell response quantified by ELISpot and depicted as spot forming units (SFU) in 100.000 PBMC (n = 20). Spots pictures are shown for a representative example of a TBEV pre-vaccinated and naïve individual. Significance compares B cell counts between groups on day 14 (F) and 28 (G) H-I) Spearman correlation between the IgG titer (H) and IgM titer (I) with the YF17D polyclonal neutralizing titer of sera at day 28. J) Neutralization curves of undepleted polyclonal serum and IgG or IgM depleted serum (in grey) for TBEV pre-vaccinated or naive individuals. K-L) Quantification of the 80% neutralization cutoff before and after IgG (K) and IgM (L) depletions. TBEV-vaccinated group is represented in orange and TBEV naïve in blue. Curve fitting in J was calculated with local regression for n = 45 TBEV vaccinated and n = 19 TBEV naive in IgG depleted serum and respective undepleted serum and n = 26 and 22 individuals per group in IgM depleted serum. Statistical significance is shown between undepleted and depleted samples using a paired Wilcoxon Matched pairs test and between TBEV pre-vaccination status using the unpaired Mann Whitney test. *P<0.05, **P<0.01,***P<0.001, ****P<0.0001 (paired Wilcoxon Matched pairs test for D, K and L and unpaired Mann Whitney Test for A, B, D, F, G, K and L).

The presence of YF17D virion-specific IgM in serum was measured longitudinally by ELISA. The IgM titer reached a plateau between day 14 and 28 pv and was comparable in both groups of vaccinees as confirmed in IgG depleted serum samples (Fig. 2C, 2D). Unlike the IgM response, participants with a prior TBEV exposure had YF17D cross-reactive IgG antibodies already at baseline and, upon vaccination, the IgG titer was further boosted resulting in a 100-fold higher titer compared to the flavivirus naïve group. The same dynamic was observed for E protein (Fig. 2E) and full-virion specific IgG (Extended Data Fig. 1A-B). The IgG titer continued to increase from day 14 to day 28 pv, at which timepoint all the study participants had seroconverted. While TBEV pre-vaccinated donors showed an anti-E IgG response already at day 14 pv, only a fraction of the TBEV naïve participants generated detectable anti-E IgG levels at that timepoint, suggesting an earlier response to YF17D in individuals with immune experience (Fig. 2E).

To assess the generation of vaccine-specific B cells, we implemented a soluble (s)E-specific ELISpot assay. The number of plasmablasts was quantified directly *ex vivo* and the total number of sE-specific B cells was measured following the differentiation of B cells into antibody-secreting cells. sE-specific IgG secreting plasmablasts peaked at day 14 pv in TBEV vaccinated participants but were below detection for naïve donors (Fig. 2F). The total amount of sE-specific B cells was in line with the IgG levels measured in serum with a significantly higher B cell number in flavivirus experienced individuals. Likewise, sE-specific B cells were detected earlier, on day 14, in TBEV pre-immunized individuals (Fig. 2G).

Collectively, these results indicate that TBEV pre-immunization does not hinder the response to YF17D. TBEV-immunized individuals have cross-reactive IgG antibodies to YF17D and experience an earlier and stronger IgG response.

### Results 2. IgM antibodies account for most of the neutralizing capacity at day 14 and 28 post immunization

The observed differences in the IgG titer did not correspond with the similar neutralization capacity between both groups of vaccinees. When groups were analyzed separately, the IgG titer at day 28 correlated weakly with neutralization (R = 0.29, p 0.04 and R = 0.34, p < 0.0001), whereas there was a stronger association with the IgM levels (R = 0.55, p < 0.0001 and R = 0.64, p < 0.0001 for TBEV naïve and experienced individuals, respectively) (Fig. 2H-I). To precisely define the contribution of IgG and IgM antibodies to neutralization we depleted serum of the IgM or IgG fractions in a subgroup of the study cohort. We confirmed that depletions were successful, without loss of the alternative antibody fraction (Extended Data Fig.1 C). We observed that the IgM fraction was the main mediator of virus neutralization, accounting for approximately 75% of the neutralizing titer on day 28 (Fig. 2J-L). Consistently, IgG depletion led only to a 25% loss of neutralization capacity on day 14 and 28 (Fig. 2J, K and Extended Data Fig. 1D-E). IgM depleted sera of TBEV-immunized individuals showed a significantly higher neutralizing capacity, indicating that the increased IgG titer contribute partly to virus neutralization (Fig. 2L). There was no significant difference in the IgM antibody titers and IgM-mediated neutralization between groups (Fig. 2D, J-K).

Thus, on day 28 the IgM fraction is primarily responsible for the neutralization capacity and is equally strong regardless of the pre-vaccination status. The IgG fraction can mediate neutralization, but the boosted response in TBEV-experienced individuals is predominantly directed towards poor-neutralizing epitopes.

### Results 3. TBEV-induced immunity mediates ADE of YF17D virus infection

As reported by Chan et al. 2016, cross-reactive non-neutralizing or sub-neutralizing IgG antibodies can mediate FcγR engagement and increase vaccine immunogenicity via ADE (21,22). Given that TBEV vaccinated individuals had YFV cross-reactive antibodies at baseline, we measured whether they could facilitate YF17D infection of FcγR expressing cell lines (THP-1 and K562). We observed an enhanced infection of a venus-fluorescent YF17D virus in both cell lines in presence of serum from TBEV-immunized donors. ADE is IgG dependent as it was absent in IgG depleted serum but not in IgM depleted serum and it was inhibited in presence of FcγR-blocking antibodies (Fig 3A and Extended Data Fig. 2 A, B). YF17D infection enhancement was observed exclusively with sera from TBEV-experienced individuals (Fig. 3B) which showed the highest enhancing titer, calculated as area under the curve (AUC) (Fig. 3C).

**Figure 3.**
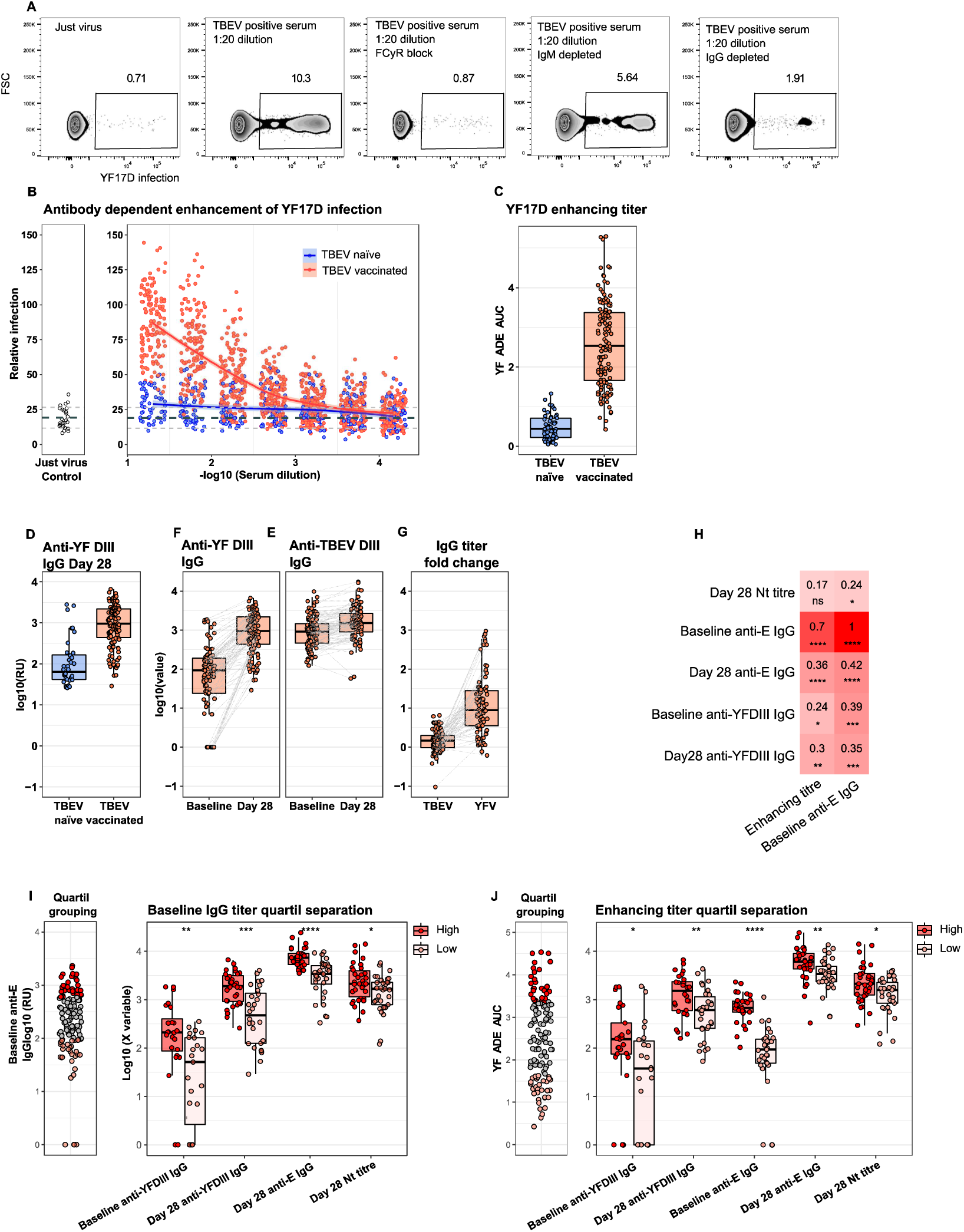
Antibody dependent enhancement of YF17D virus infection mediated by TBEV vaccine-induced IgG. A) Flow cytometric determination of venus-YF17D virus infection of THP-1 cells in absence or presence of cross-reactive serum from a TBEV-vaccinated individual. The conditions tested include polyclonal serum alone or in combination of FCyR-blocking antibodies and IgM or IgG depleted serum. B) ADE of YF17D mediated by study participant‘s serum. Virus infection was quantified in combination with serially diluted serum and was normalized against the enhancement of a 1:20 diluted cross-reactive serum of an internal control carried for all measurements. Thick dashed line indicates the mean of the different just virus controls and dotted lines define +/-1 SD. C) Quantification of the enhancing titer as AUC of the normalized virus infection across serial dilutions shown in B. D) Comparison of anti-YF17D-DIII specific IgG titer at day 28 post vaccination for both TBEV groups (n = 36 TBEV naive, n = 117 TBEV vaccinated individuals) F,E) Longitudinal (day 0 and day 28 pv) quantification of anti-TBE-DIII n = 114 pairs (F) and anti-YF17D-DIII n = 97 pairs (E) specific IgG in TBEV experienced individuals. G) Paired comparison of the DIII-specific IgG fold-change between day 28 and day 0 for TBEV and YF17D. H) Spearman correlation in TBEV experienced individuals of baseline anti-E IgG and enhancing titers versus YF17D vaccine-induced neutralizing antibody titer at day 28, anti-sE and anti-DIII IgG titers and baseline anti-YF17D-E and anti-YF17D-DIII IgG titers. Color intensity and number reflects the spearman correlation coefficient. I,J) Comparison of the first and fourth quartile group of YF17D-sE specific IgG (I) and Enhancing titers (J) at baseline and distinct parameters of the humoral response. P<0.05, **P<0.01, ***P<0.001,****P<0.0001 (unpaired Mann Whitney Test for I-J).

In addition, we quantified the IgG fraction targeting DIII. DIII is often used for serological diagnosis (38) and, although cross-reactive epitopes have been described (39,40), responses to DIII are generally virus-specific (41). As observed for anti-sE IgG antibodies, TBEV pre-vaccinated individuals had a stronger IgG response against YF17D-DIII (Fig. 3D). We then compared the fold change in anti-YF17D-DIII and anti-TBEV-DIII IgG titers between baseline and day 28 pv. Interestingly, the strong expansion of anti-YF17D-DIII-reactive IgG (10-100-fold) contrasted with the moderate increase in TBEV-DIII specific IgG (<10-fold) (Fig. 3F-E). Since the IgG fraction targeting TBEV-DIII was not expanded to the same extent, we conclude that YF17D is not only potentially boosting cross-reactive TBEV-induced memory responses, but via an enhanced immunogenicity also triggers antibodies towards previously unseen, non-cross-reactive epitopes in DIII.

The correlation between the baseline levels of sE-specific IgG and the baseline enhancing titer with the post-immunization IgG response to sE and DIII, as well as the post-vaccination neutralizing titers, suggests that ADE mediates an increase in vaccine immunogenicity (Fig. 3H and Extended Data Fig. 2C-E). To gain insight into the size effects of ADE on vaccine immunogenicity, we grouped the vaccinees into quartiles based on their enhancing and baseline IgG titers (see methods section). This approach removes from the analysis “average” enhancers or individuals with intermediate TBEV-vaccine-induced IgG levels and therefore improves precise identification of true differences between the high and low vaccine enhancers.

Participants in the highest quartile of the enhancing titer had a significantly higher neutralizing titer and IgG levels against sE and DIII (Fig. 3I) than the participants in the lowest quartile. The same associations were observed with baseline IgG quartiles (Fig. 3J). Thus, within the TBEV-pre-vaccinated group, ADE of YF17D was associated with increased vaccine immunogenicity.

Altogether, these results suggest that TBEV pre-immunity may increase the magnitude and breadth of the YF17D vaccine response.

### *Results 4.* YF17D effectively primes for non-cross-reactive antibodies in flavivirus-naïve vaccinees but expands a pan-flavivirus cross-reactive IgG response in TBEV-experienced individuals

The IgG response in TBEV-experienced and naïve vaccinees was investigated for cross-reactivity with other members of the *Flaviviridae* family. Using an indirect immunofluorescence test, the IgG and IgM binding to ZIKV, JEV, WNV, TBE, YFV and all serotypes of DENV was measured in a cohort subset (Fig. 4 and Extended Data Fig.3). At baseline, TBEV vaccination induced IgG cross-reacting with all flaviviruses at similar magnitude. This pan-flavivirus cross-reactive IgG signature was boosted upon vaccination with the YF17D vaccine. In contrast, the YF17D vaccine induced an IgG response that targeted uniquely YFV in naïve individuals (Fig. 4A). Consistent with previous studies, the IgM signature was YFV-specific and could not be detected at baseline (Fig. 4B).

**Figure 4.**
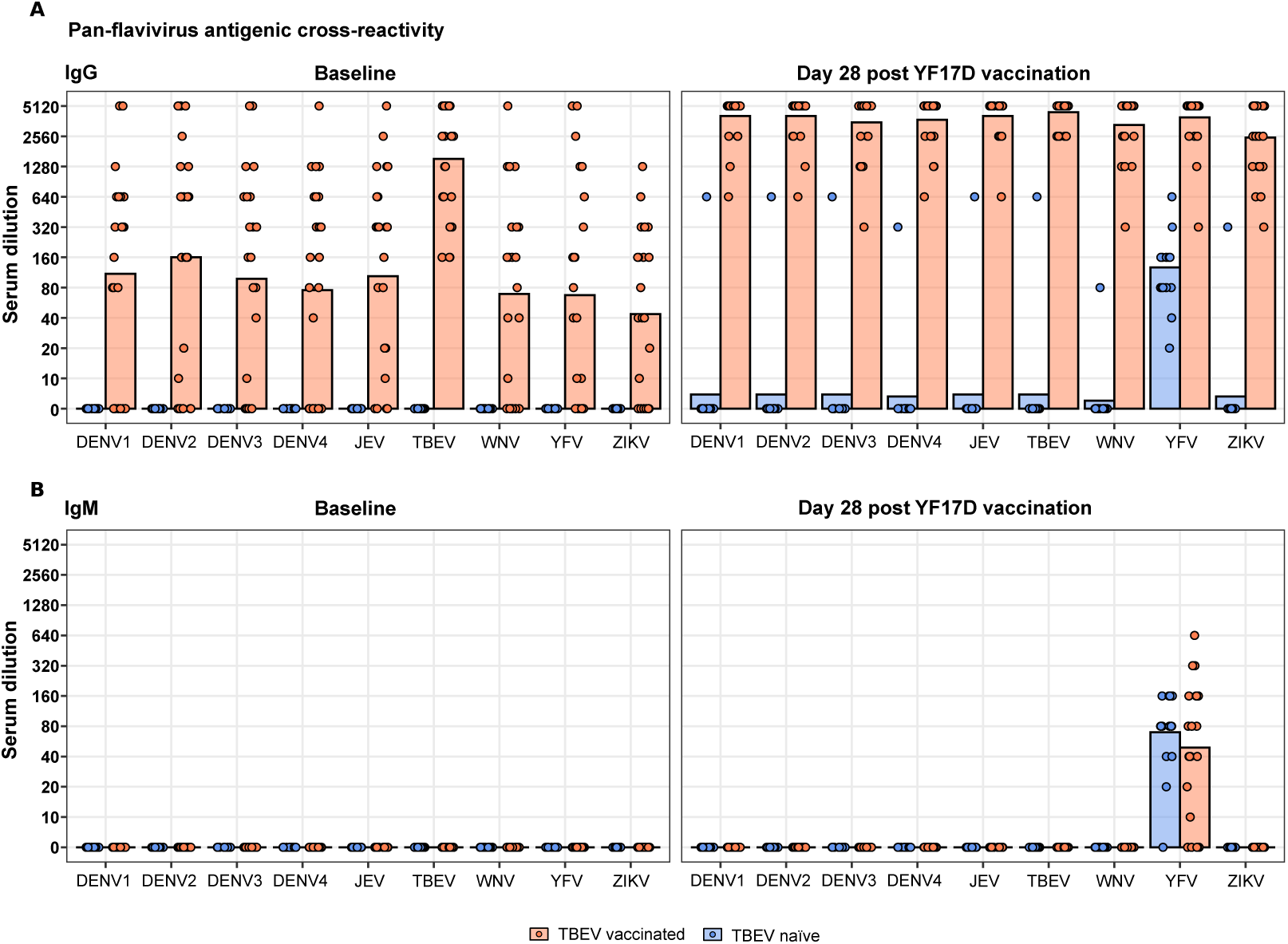
IgG and IgM cross-reactivity signature before and after YF17D vaccination in TBEV naïve and pre-vaccinated individuals. A-B) IgG (A) and IgM (B) subtype cross-reactivity evaluation before and after YF17D vaccination using an indirect immunofluorescent test for a panel of nine human-pathogenic flaviviruses: DENV 1-4, ZIKV, WNV, JEV, YFV and TBEV. A subgroup of n = 39 individuals was tested (Extended data Fig. 3), out of which n = 15 were TBEV naïve and n = 24 TBEV experienced. Bars indicate titer mean and dots reflect the antibody amounts as serum dilution end-point titers

These results highlight the capacity of the vaccine strain of YFV to prevent a cross-reactive response when administered to flavivirus naïve individuals.

### Results 5. Pre-existing immunity changes YF17D immunodominance and misdirects the response to FLE

Depending on previous flavivirus exposure, YF17D induces a differential cross-reactivity pattern while eliciting a comparable neutralizing response. We hypothesized that the neutralizing capacity is predominantly driven by antibodies targeting EDE and the FL-proximal region whereas cross-reactive antibodies aim at the immunodominant FLE. To better understand this change in the immunodominance we designed a set of recombinant sE protein mutants for the study of the IgG response to different epitopes (Fig. 5A). Besides the monomeric sE protein containing all three ectodomains and variants consisting of either only DI-II or only DIII, we designed constructs displaying quaternary dimeric epitopes to reproduce the epitope landscape of YF17D more realistically. The substitution S253C in DII allows the formation of an inter-protomer disulphide bond across the two sE protomers generating a covalently bound dimer (42). This construct (hereinafter referred to as breathing-dimer) retains the ability to oscillate and exposes EDE, FLE and the FL-proximal region. Furthermore, a W101H substitution was introduced in the breathing-dimer setting to disrupt the FLE (breathing-dimer^W101H^) (Fig.5 A). In addition, we produced a locked-dimer E protein by introducing a double substitution L107C and T311C following the strategy used by Rouvinski et al (2017) and Slon-Campos et al (2019) with DENV and ZIKV. This construct displays quaternary epitopes on a pre-fusion dimeric structure that is bridged with two disulfide bonds between DI and DIII of opposing protomer units (Extended Data Fig. 4A-B).

**Figure 5.**
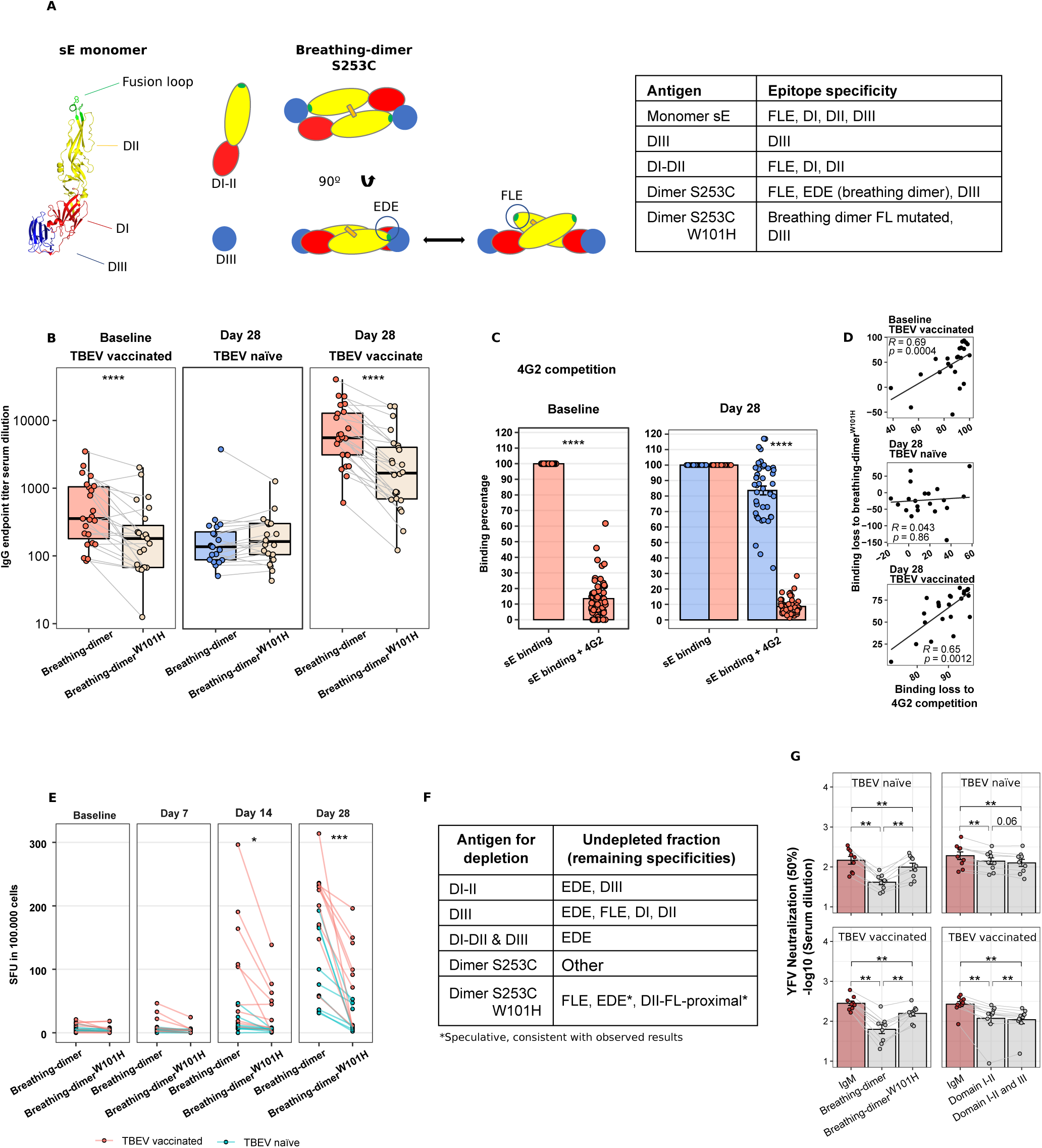
Binding sites and neutralizing capacity of the YF17D vaccine-induced IgG response in TBEV naïve and pre-vaccinated individuals. A) Recombinant proteins for the dissection and functional analysis of different antibody specificities. Illustration depicts the envelope protein ectodomains (sE), DI-II, and DIII produced separately as well as recombinantly produced dimeric structures. The table summarizes the epitopes displayed by the protein antigens used. B) IgG endpoint titer quantification for breathing-dimer and breathing-dimer^W101H^ specificities by ELISA for both TBEV groups at baseline and Day 28. C) Antibody binding competition to sE of participants serum with the FL-mab 4G2. Percentage of remaining binding is calculated by comparing the binding signal with and without 4G2 as competitor D) Spearman correlation between antibody binding loss estimated with the 4G2 competition assay (C) and with the breathing-dimer^W101H^ (B). E) Longitudinal quantification of IgG-producing B-cells specific for breathing-dimer and breathing-dimer^W101H^. Units represent spot-forming units per 100.000 PBMC. F) Table describing the antigens used for antigen-specific IgG depletions and the expected specificities of the remaining undepleted faction used for YF17D neutralization assays. G) YF17D neutralization titers (50% cutoff) of IgM depleted and antigen-specific depleted sera as explained in F. Envelope structure accession number (PDB: 6IW4) was edited using Pymol. For the ELISpot assays n = 20. For antibody titer quantification a subgroup of the study cohort was used (TBEV vaccinated = 24, TBEV naïve = 24). For antigen depletions n=20 (TBEV vaccinated = 10, TBEV naïve = 10). Individual selection is shown in Extended Data Fig. 3. *P<0.05, **P<0.01,***P<0.001, ****P<0.0001 (paired Wilcoxon Matched pairs test except for C, for which unpaired Mann Whitney Test was used).

The comparison between the breathing-dimer and breathing-dimer^W101H^-specific IgG titers serves to measure the fraction targeting FLE. For TBEV-experienced individuals, the antibody fraction targeting the breathing-dimer was significantly reduced in baseline samples and at day 28 by the W101H mutation (45 and 64% reduction respectively) whereas TBEV naïve individuals showed no significant difference (Fig. 5B). Complementary, the sE-specific IgG titer was quantified in binding competition assays with 4G2 (pan-flavivirus FLE-specific mAb) and 2D12 (YFV-neutralizing, non-cross-reactive anti-E mAb) (Fig. 5 C and Extended Data Fig. 5A). As anticipated, TBEV pre-exposed vaccinees lost over 80% of the YF-IgG binding fraction at day 28 in competition with 4G2 while flavivirus naïve individuals ranged from 0 to 60% binding loss demonstrating that FLE is a dominant binding site for the antibody response in TBEV-experienced individuals, but not in TBEV-naïve individuals. Likewise, baseline antibodies also competed with 4G2 for binding (Fig. 5C). Consistently, binding loss caused by W101H mutation and competition with 4G2 correlated with each other (R = 0.65, p = 0.0012), serving as cross-validation of these assays to quantify FLE antibodies in serum (Fig. 5D).

Additionally, we performed an ELISpot assay to quantify the number of epitope-specific circulating B cells. We observed that the number of breathing-dimer-specific B cells was larger for TBEV pre-immunized compared to TBEV-naive participants. Approximately 50% of the specific-B cells in TBEV pre-immunized donors (100 cells/100.000 lymphocytes) required the unmutated FL for binding (Fig. 5E). Consistently with serum antibody levels, TBEV naïve individuals had lower numbers of breathing-dimer specific B cells, although a relevant fraction produced antibodies requiring FL for binding. These B cells release antibodies that may be binding dimeric structures or FL-proximal regions whose binding site includes aminoacids located in the FL (Fig. 5A and E). The locked-dimer-specific IgG and B cell response was in line with our previous findings showing increased responses in TBE-experienced individuals (Extended Data Fig. 4 C-D).

Taken together, these results show that the IgG fraction targeting the FLE is dominant in TBEV pre-immunized but not in naive participants. However, the fusion loop region is a binding site for antibodies elicited in both groups.

### Results 6. Antibodies targeting quaternary E-protein epitopes are central for neutralization of YFV in TBEV-experienced and naïve vaccinees

To assess the neutralizing capacity of antibodies with different specificities, we performed antigen-specific IgG depletion from serum samples that had been pre-depleted of IgM antibodies (Methods and Fig. 5F-G). By depleting the IgM fraction, the main neutralizing sites targeted by the long-term, durable, IgG response can be more precisely dissected. As expected, IgM removal reduced greatly the neutralizing capacity of the sera (Fig. 2G). Further depletion with the breathing-dimer protein resulted in a remarkable loss of neutralizing capacity to a similar extent in TBEV-experienced and naive individuals. The neutralizing titers of sera depleted with the breathing-dimer^W101H^ antigen remained high, proving that neutralizing epitopes include the FL as a binding site (Fig. 5 G, left panels). In fact, the main chain of the FL is part of the EDE epitope (10). Depletions performed with the locked-dimer construct resulted only in a slight decrease in neutralizing capacity (Extended Data Fig. 5E). Given the occlusion of the FL in the locked dimer we thought this construct would deplete the EDE antibodies and would reduce greatly the serum neutralization activity. However, the direct modification of the FLE with the L107C mutation of this construct had also an impact on the EDE binding site, affecting the ability to deplete these antibodies. A comparable construct for dengue (43) although able to bind most of the dengue EDE antibodies showed reduction in binding for selected EDE. Similarly, the YFV E locked-dimer construct may have failed to deplete the principal antibody fraction responsible for the virus neutralization (Extended Data Fig. 4A-E).

sE monomer cannot be used to deplete exclusively monomer-specific IgG antibodies since EDE antibodies may assemble sE monomers together into dimers and therefore, this construct would deplete also antibodies with dimer-specificities (44). To ensure the removal of antibodies with monomeric but not dimeric specificities, subsequent depletions with DI-II and DIII were then performed. Even though this resulted in a clear loss of neutralizing capacity, especially in TBEV-experienced individuals, monomeric specificities only made up a minor fraction of the polyclonal neutralizing antibody response when compared to the breathing-dimer depleted sera (Fig. 5G right panels).

Altogether, these results highlight the importance of EDE as the main neutralizing site and reveal that the FL is a critical component of the binding site for potent neutralizing EDE antibodies.

### Results 7. YF17D vaccination boosts antibody dependent enhancement of DENV and ZIKV in TBEV pre-immunized individuals driven by FLE antibodies

Given that in TBEV-experienced individuals YF17D boosts a pan-flavivirus cross-reactive IgG response, we examined whether these individuals mediate ADE to dengue and Zika viruses using viral reporter replicon particles (VRP) (45).

Interestingly, the antibody response induced by TBEV immunization was sufficient to enhance DENV and ZIKV infection. The enhancing capacity was further increased after YF17D vaccination. As expected, flavivirus naïve individuals did not facilitate DENV and ZIKV infection at baseline and, importantly, the YF17D vaccine did not induce antibodies with enhancing potential. This is consistent with the absence of cross-reactive antibodies in these individuals (Fig. 6A).

**Figure 6.**
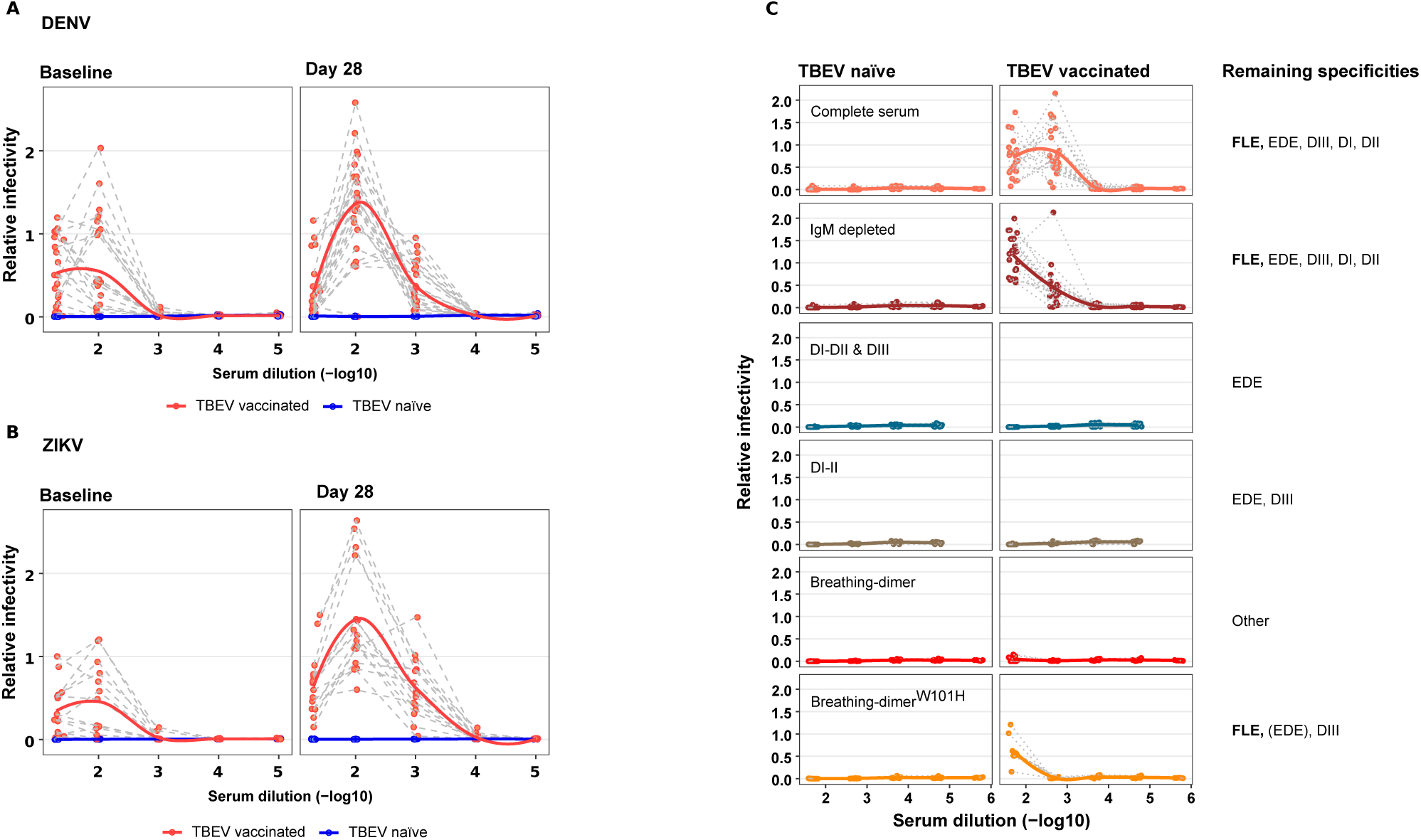
Antibody dependent enhancement of DENV and ZIKV by bulk and antigen-specific IgG depleted serum in TBEV naïve and pre-immunized individuals before and after YF17D vaccination. A) Antibody dependent enhancement of dengue and Zika VRPs of TBEV-experienced and naïve sera at baseline and day 28 post-YF17D vaccination. Dengue ADE n = 40 (20 per group), ZIKV ADE n = 24 (12 per group). B) Dengue ADE driven by: Undepleted, IgM depleted and IgM & antigen-specific-IgG depleted sera (as figure 5 F) Depleted serum was tested for 20 individuals (10 per group). Relative infectivity is estimated as the normalized fold-increase of infection to an internal control carried for all the assays.

To elucidate the antibody specificities mediating ADE, we first removed the IgM fraction and then measured the enhancing capacity after antigen-specific depletions of the remaining IgG fraction. ADE to dengue virus was lost in serum depleted of antibodies binding to the breathing-dimer, sE monomer (DI-II and III) or DI-II. In contrast, samples depleted with breathing-dimer^W101H^ or locked-dimer constructs retained antibodies mediating ADE (Fig. 6B and Extended Data Fig. 5F). These results point to FLE-antibodies as responsible for cross-reactivity and ADE.

In conclusion, YF17D vaccine expands FLE-antibodies with potential to mediate enhanced dengue and Zika infection in TBEV pre-immunized donors. Nevertheless, in a flavivirus naïve population, YF17D primes for non-cross-reactive but effectively neutralizing antibodies.

## Discussion

Circulating and emerging flavivirus infections represent a major health concern. The reduction of their burden and impact requires vaccination strategies based on a deep understanding of anti-flavivirus immunity. YF17D was empirically developed (27) and achieved an optimal balance between immunogenicity and attenuation. Some studies have addressed the reasons for the outstanding YF17D vaccine performance (46–48) but the immunological basis for its effectiveness remains poorly defined. The global distribution and high prevalence of flaviviruses create the need to understand and integrate the effect of pre-existing immunity to antigenically related flaviviruses in vaccine design and other anti-flavivirus strategies.

In this study, we have characterized the antibody response elicited by the YF17D vaccine in a cohort of individuals who were either TBEV pre-immunized or unexperienced. By day 28 post vaccination, an adaptive response had been mounted as shown by a 100% seroconversion rate and protective neutralization titers in the study participants. At that timepoint, TBEV pre-immunity did not influence the polyclonal neutralizing response, which was largely dependent on IgM. The predominant IgM contribution to flavivirus neutralization may be achieved through its multivalent binding to the multimeric epitope arrangement of the virion and can persist for several years after infection or vaccination (49,50). Nevertheless, by day 28, the IgG fraction already contributed to neutralization with higher efficacy in TBEV pre-immunized individuals. The long-term effects of pre-existing immunity on YF17D-induced neutralization, cross-reactivity and protection durability have not been addressed in this study. We speculate that while the IgM fraction gradually wanes, the IgG compartment will persist and partake most of the neutralization activity. Therefore, differences in neutralization titers across both groups may become more apparent at later time points.

YF17D virus seems to be concealing cross-reactive epitopes while priming for the generation of neutralizing antibodies. This capacity is not observed in other flavivirus infections or vaccinations which inevitably generate cross-reactive antibodies (10,35,51). We hypothesize that the specific structure of the YF17D virion leads to limited exposure of the FLE. Thus, the formation of neutralizing antibodies targeting complex quaternary epitopes as well as the FL-proximal region is structurally favored. This particular priming might explain the lack of association between YF17D vaccination and dengue severity (52) contrary to the JEV vaccine which induces a cross-reactive response and is associated with severe dengue disease (19,53).

An extensive body of literature has shed light on the relevance of the FL-proximal site for YFV neutralization. This conserved area in DII contains epitopes targeted by well-characterized protective neutralizing antibodies (29–32). In our study, we show that quaternary epitope binders constitute a substantial fraction of the neutralizing activity of the polyclonal human sera post YF17D vaccination. Even though EDE is a known neutralizing site in flavivirus immunity (10,11), up to date, a precise mapping of EDE antibodies for YFV has not been published. Besides, their characterization could not be done in previous studies when monomeric sE was used for B-cell sorting (31,32). The relevance of EDE-antibodies in anti-YFV immunity was hinted before. Monomeric specificities did not completely account for the neutralizing activity in polyclonal human sera (33) and the generation of viral escape variants to potent neutralizing antibodies mapped sites compatible with an EDE in YFV (28). To gain insight into these complex quaternary epitopes we used covalently bound dimers with potential to display EDE. We observed a profound loss of neutralizing activity upon removal of dimer-specific IgG while sera depleted of IgGs targeting DI-II and DIII largely preserved the capacity to neutralize YF17D. The dissection of antibody specificities in polyclonal serum is challenging given that EDE, FLE and FL-proximal epitopes overlap in some positions in the fusion loop. Nevertheless, our toolbox proved useful for the dissection of the differential antibody specificities elicited upon YF17D vaccination and our results highlight dimer epitopes as key determinants of YF17D neutralization.

A clear differential pattern in the magnitude, breadth, and specificities of the IgG response to YF17D distinguished flavivirus experienced from unexperienced individuals. Despite the finely tuned epitope landscape of the YF17D vaccine, the antibody response is skewed towards the FLE in TBEV-experienced individuals. We cannot rule out the possibility of an immune imprinting conferred by the TBEV vaccine which is now dominating the response to YF17D. Nevertheless, we showed that the breadth of the humoral response cannot be explained exclusively by a recall response to conserved epitopes. We, along with others, observe a robust cross-reactive antibody response induced by the TBEV vaccines (54) which share antigenic determinants with YF17D virus including the FLE (except for a G104H change in the FL sequence). Upon heterologous vaccinations, cross-reactive clones of the memory B cell (MBC) compartment can dominate over *de novo* or mature B cells undergoing germinal center reactions and hence restrict the final diversity of the B cell response to a new challenge (55). MBC recognition of the epitope is a prerequisite to initiate a recall response but the concealment of the cross-reactive FLE by the YF17D virion would hinder this process. Nevertheless, FLE might still be transiently exposed and FL-Ab may be induced in low amounts (32,33) although minimal cross-reactivity following vaccination is observed (37,56). The FL-dominant response in TBEV pre-vaccinated individuals may result from a recall response by MBC to a reduced but sufficiently exposed FLE. Alternatively, FLE accessibility could be modulated upon binding of cross-reactive antibodies to the YF17D virion favoring FLE dominance.

We demonstrated that TBEV-induced cross-reactive antibodies can enhance YF17D infection via ADE. In addition, we show that higher enhancing titers associated with stronger neutralizing antibody responses suggesting increased vaccine immunogenicity. This is consistent with a previous study by Chan et al (2019) which showed that JEV vaccine-induced cross-reactive antibodies enhance YF17D immunogenicity via ADE (21).

The efficacy of both TBEV and YF17D vaccines is excellent (26,57). We show that vaccination against TBEV followed by YF17D does not hamper the protective capacity of these vaccines. However, when administered in reverse order, YF17D vaccinated individuals were shown to generate comparable IgG titers, with increased cross-reactivity but poorer neutralizing capacity to TBEV (34,36). Here we report that sequential flavivirus exposures increase antibody cross-reactivity and enhance infection by heterologous flaviviruses. ADE is clinically relevant in heterotypic secondary DENV infections (16,17) and JEV vaccine responses have been clinically associated with dengue severity (19). Likewise, we infer that sequential TBEV and YF17D vaccinations could render these individuals susceptible to severe dengue and Zika disease. The double exposure to TBEV and YF17D vaccine is not uncommon among European and Asian travelers living in TBEV endemic areas. In fact, dengue has become one of the most important emerging diseases among European travelers (58) with an increasing incidence in recent years (59,60). The predisposition to severe dengue disease should be considered for awareness and personal prevention measures until unequivocal epidemiological data confirms or rejects this hypothesis.

In summary, our data demonstrates that the immunogenicity and antibody immunodominance of a live vaccine is not only influenced by structural factors of the immunogen and the biology of the vaccine virus but also by the pre-exposure to related antigens. Pre-existing immunity governs the selection of B cell clones for antibody production balancing a response that could be neutralizing or misdirected to non-neutralizing and cross-reactive epitopes. This must be considered in structure-based vaccine designs, especially in the context of prevalent co-circulating and related pathogens, to successfully achieve responses to protective antigenic sites with minimal cross-reactivity.

## Materials and Methods

### Study samples and personal information

250 healthy young adults, naive to natural flavivirus infection as well as a negative vaccination history to JEV and YFV, were recruited from 2015-2019 at the Division of Infectious Diseases and Tropical Medicine (DIDTM) as well as the Department of Clinical Pharmacology, University Hospital, LMU Munich, Germany. The study protocol was approved by the Institutional Review Board of the Medical Faculty of LMU Munich (IRB #86-16) and adhered to the most recent version of the declaration of Helsinki. 67.6% of the study participants were females (n=169) and the median age is 24 years (range 19-47) (Fig. 1C). The body mass index fell in the normal range (18.5 – 24.9) for most participants (n=212; 84.8%) with some outliers (n=38; range 17.7-46.4). After giving informed consent, longitudinal samples were collected from venous blood draws prior to and on days 3, 7, 14 and 28 after subcutaneous YF17D vaccination (0.5 mL of Stamaril; Sanofi Pasteur, Lyon, France). Samples were analyzed retrospectively and in an anonymized manner.

#### Human blood samples

Blood samples were collected via venipuncture and serum and plasma samples were stored in aliquots at −80°C. PBMC samples were isolated manually from buffy coat following Ficoll-Paque PLUS (GE Healthcare, Sweden) density centrifugation and cryopreserved in heat-inactivated foetal calf serum (FCS) supplemented with 10% DMSO (Sigma-Aldrich) in liquid nitrogen. All cellular assays started with the thawing of cryopreserved PBMC with an average recovery of 70 %.

#### Protein cloning, production and purification

The YF17D Envelope protein ectodomains, subdomains and covalently bound dimers used in this study were produced recombinantly in stably transfected drosophila S2 cell lines (ThermoFisher Scientific, R690-07). Site directed mutagenesis (QuikChange II, Agilent, 200523) was performed to introduce a cysteine in position 253 or 107 and 311 to form the covalently bound dimers and a histidine in position 101 to disrupt FLE using the following primers and validated by DNA sequencing.

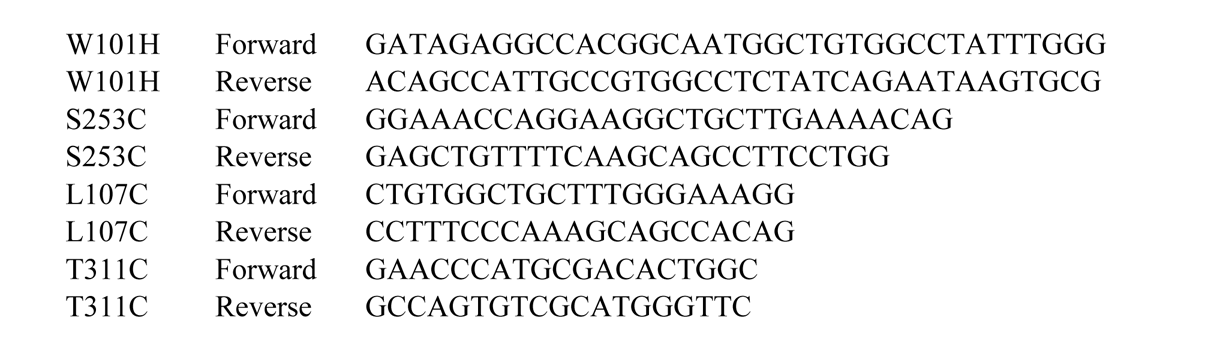

For protein production a pT350 plasmid encoding for the construct of interest with a BiP signal sequence in the N-terminal and a double strep-tag in the C-terminal region was co-transfected with a puromycin resistant plasmid (pCoPuro) (61) with effectene reagent (Qiagen, 301425) following manufacturer’s instructions. Cells were then subjected to puromycin selection to generate a polyclonal stable cell line. Protein expression was induced with 5µM CdCl_2_ in Insect-XPRESS medium (Lonza) for 5-7 days and supernatants were concentrated and purified using strep-tacting affinity chromatography columns in an AKTA FPLC system. To separate properly folded protein from aggregates, the protein was further purified by size exclusion chromatography and the elution peaks were validated by SDS-PAGE. Finally, the protein was stored in a NaCl 150mM, Tris-HCL 10mM pH 8 solution.

Protein production and purification were validated by SDS-PAGE/Western blot under reducing and non-reducing conditions. Proper folding and epitope display was confirmed upon binding of monoclonal antibodies. 5A mAb binds the FL-proximal region, the pr binding site (30) and 4G2 mAb binds the FLE. W101H successfully prevents the binding of 4G2 and reduces binding of 5A. 4G2 and 5A bind both to breathing-dimer and sE monomer but are unable to bind the locked-dimer (Extended data Fig 5B). Given the occlusion of the FL in the locked dimer setting, this construct does not permit the binding of antibodies targeting FLE and FL-proximal region.

#### YF17D and YF17D-Venus virus production

The YF17D virus was directly amplified from a Stamaril vaccine dose. YF17D variant YF17D-Venus plasmid was a generous gift from Charles M. Rice and Margaret MacDonald (The Rockefeller University, New York, USA). Virus stock production and purification was done as previously described (62,63) with small modifications. Briefly, Vero B4 cells (DMEM 10%, 2 mM L-glutamine, and 1 % pen/strep) at 37 °C and 5 % CO_2_, were expanded and infected at an MOI 0.1. Supernatants were collected post-infection when a clear cytopathic effect was visible. Supernatants were then combined with polyethylene glycol (PEG 8000) 7% (w/v) and pellet was resuspended and homogenised for final purification by sucrose cushion separation after ultracentrifugation in an MLS 50 rotor (Beckman). The final purified virus stock was diluted in TNE buffer (20 mM Tris-HCl pH 8, 150 mM NaCl, 2 mM EDTA) quantified by plaque assay and stored at −80°C until use.

#### TBEV plaque reduction neutralization tests (PRNT)

A549 lung adenocarcinoma cells (ATCC® CCL-185™) were cultured as confluent monolayer (MEM, 1x NEAA, 10 % FBS (Gibco, Thermo Fisher Scientific Inc.) at 37 °C and 5 % CO2. The TBEV reference strain Neudoerfl was used and initially passaged twice on A549 cells (MEM, 1x NEAA, 2 % FBS, 4 days) to provide a sufficient virus stock solution (MEM, 1x NEAA,20 % FBS), which was subsequently kept at −80 °C. The virus titer of the stock solution was determined by performing a PRNT with ten-fold dilutions and calculated according to Baer & Kehn-Hall 2014 (64). Confluent monolayers of A549 cells were grown in 24-well/96-well plates for 24 hours. The plasma samples were inactivated at 56 °C for 30 min and a series of seven dilutions (1:5 – 1:320) were prepared. TBEV, diluted to a concentration of 500 pfu/ml, was mixed in equal volumes with the plasma dilutions and incubated for one hour at 37 °C and 5 % CO_2_. Next, cells were washed with PBS and inoculated with 100/20 µL of virus-plasma mixture in triplicates. Positive and negative controls were also added in triplicates on each plate. Plates were incubated for one hour at 37 °C and 5 % CO_2_, subsequently washed with PBS and 500/100 µL CMC medium (MEM, 1x NEAA, 2% FCS, 0.75 % carboxymethylcellulose) was added and incubated for three days at 37 °C and 5 % CO_2_. Following supernatant removal, plates were stained with crystal violet (13% formaldehyde, 0.1 % crystal violet) at 4 °C, washed and dried for visual examination. The neutralization antibody titer was calculated as mean plaque count of the positive control and was multiplied by 0.1 and 0.2 to obtain the cut-off value that implies a plaque reduction of 90 and 80 %. The mean plaque count of every dilution triplicate was determined and compared with the cut-off value. The first plasma dilution that resulted in a mean plaque count below the cut-off value, was interpreted as to be the neutralization titer of the plasma sample.

#### Indirect immunofluorescence tests (IIFT)

Plasma samples of 39 study participants were tested for IgM as well as IgG reactive to TBEV, WNV, JEV, YFV, DENV (types 1-4) by using EUROIMMUN Flavivirus Mosaic 3 and ZIKV IIFT assays (EUROIMMUN Medizinische Labordiagnostika AG, Lübeck, Germany) according to the manufacturer’s instructions. For standardized analyses, the titerplane technique, as described in Niedrig et al. 2008 (65), and the automated washing system MERGITE! (Euroimmun AG, Lübeck, Germany) was used. For antigen-specific IgM detection, samples were prepared with a RF absorbent (EUROSORB, 2% Tween). After incubation of biochips with plasma samples, FITC-labeled anti-human IgM or IgG was added. The MERGITE! washing step with PBS-Tween pH 7.2 was performed after each incubation period of 30min at room temperature. Thereafter slides were covered with glycerol and cover glass, before being examined using a fluorescent microscope (EUROStar 3 PLUS, EUROIMMUN Medizinische Labordiagnostika AG, Lübeck, Germany) at 460-480nm wavelength. Antibody titers were determined at serial two-fold dilutions from 1:10 to 1:5120 including suited controls. Any specific perinuclear fluorescence for IgM or IgG was considered as a positive reaction. Samples with titers below 1:10 were considered negative and samples with a titer of 1:5120 were considered as ≥1:5120.

#### Determination of YF17D neutralizing antibody titers

The neutralizing antibody titer was determined by a Fluorescence Reduction Neutralization Test (FluoRNT) as previously described by Scheck et al, 2022 (63). Briefly, YF17D-Venus virus, was incubated with equal amounts of heat-inactivated and serially diluted donor sera for 1 h at 37°C in serum-free DMEM. 21.000 PFU/well of virus was used to reach approximately a 50% cell infection. The mixture was then added to Vero cells (25.000 cells/well in a 96-well plate) and incubated for 24h. Cells were then trypsinised, stained for viability (ThermoFischer, L10120) and fixed with 4% PFA before acquisition in FACS Canto or CytoFLEX LX. Frequency of infected cells in the absence of vaccinee serum was set as 100% and the percentage of reduction was calculated for each dilution step as follows.

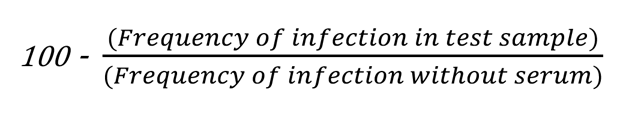

Neutralization curves were fitted by 4 parameter logistic regression using Prism 8 (GraphPad, La Jolla, CA, USA) or drm function of drc R package. 50, 80 and 90 % FluoRNT values were interpolated from the curves.

#### Quantification of anti-YF antibodies in human serum

An in-house three-layer ELISA was used for the quantification of antigen-specific IgG and IgM in vaccinees serum. Half-area 96 well plates (corning) were coated overnight at 4℃ with 1 µg/ml or 3.5 µg/ml of sE protein or full virus YF-17D respectively (estimated by bradford assay) in 0.1 M sodium carbonate buffer pH 9.5. For determining anti-DIII IgG responses, 1 µg/ml of recombinant YF17D-DIII or TBEV-DIII (Jena Bioscience, PR-1450-S) was used. Only full virus YF17D was used as antigen for the quantification of vaccinee-specific IgM. Following washes with PBS tween-20 0.05% the plates were blocked with 10% goat serum in PBS+tween-20 0.05% for 3h at room temperature and 200 rpm shaking. Next, heat-inactivated serum samples were added in three-fold dilutions −typically 1:100, 1:300, 1:900 and 1:1800 for IgG detection and 1:200, 1:400 for IgM)-in blocking buffer and incubated for 2 h. To ensure the removal of unspecific immunoglobulins, plates were thoroughly washed before the addition of anti-human IgG-HRP (Jackson ImmunoResearch, 109-035-088, 1:5000) or anti-human IgM-HRP (ThermoFischer, 31415, 1:5000). One hour later, plates were washed again with PBS+Tween-20 0.05% followed by the addition of TMB substrate (BD, 555214). Plates were allowed to develop for 10-25 minutes before adding 1M H_2_SO_4_ stop solution. The optical density was read at 450 nm with 595 nm correction. Signal was considered positive if at least 3x-higher than background measured in wells without pre-coated antigen and same serum dilution. Background signal was subtracted. To ensure inter-day and inter-plate comparability and reproducibility, ELISA titers are reported as relative units (RU) to a standard anti-YF17D serum given an arbitrary value of 10000 units as described previously (35,66,67). The corresponding values for the test serum samples are read from the standard curve fitted using a four-parameter logistic regression.

#### Competition ELISA

To determine the amount of IgG antibodies in human serum directed to FLE, we measured the loss of antibody binding to sE protein in competition with a murine isotype of 4G2 and 2D12 monoclonal antibodies by ELISA. As described above, ELISA plates were coated with 1 µg/ml of sE protein and blocked with 10% goat serum in PBS+tween-20 0.05% and an equimolar concentration (35 nmol) of 4G2 or 2D12 mAb in corresponding wells for 3h. Samples were tested in 1:100 and 1:1000 dilutions with 35 nmol of 4G2 or 2D12. ELISA titers, reported as RU, were compared to the titer quantified in absence of any competing antibody and the amount of anti-FLE antibodies present in the sample was calculated as follows.

> % anti-FL antibodies = [1-Sample with 4G2/Sample without 4G2]*100%

#### Endpoint titer IgG ELISA

IgG antibody titers directed against locked-dimer, breathing-dimer and breathing-dimer^W101H^ antigens were quantified as endpoint titers. In brief, following coating and blocking of the ELISA plates, seven serial dilutions of the serum samples (starting 1:50 and diluted 1:3) were added to the plates. Following a 2 h incubation, plates were washed as described above with two additional washes with PBS + NaCl 650 mM. The addition of a chaotropic agent helps to remove antibodies binding with low affinity (REF) but does not affect the quantification of antigen-specific antibodies or the protein structure therefore making the comparison and quantification of antibodies targeting antigens displaying or disrupting epitopes more reliable. This way, we could more clearly delineate the W101H impact on antibody binding between breathing-dimer and breathing-dimer^W101H^. The antibody titer is reported as the interpolated dilution of donor serum where the cutoff of OD value of 0.1 after background subtraction was reached.

The frequency of FL-specific antibodies was calculated as follows

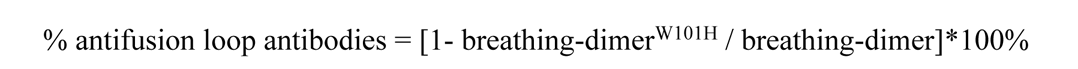

#### Bulk IgM depletion

Serum, diluted 1:3 in PBS, was added to anti-human IgM agarose beads (Sigma, A9935) for 1.5h at RT. Double volume amounts of beads were used per depletion round. Beads were pre-washed with PBS and supernatants were removed following a centrifugation (5 min 400 g). To ensure complete IgM removal, two depletion rounds were performed and verified by YF17D IgM and IgG ELISA.

#### Bulk IgG depletion

For complete removal of the IgG fraction in serum samples we used Protein G spin plates (Thermo, 45204) following the manufacturer’s instructions with minor modifications. Briefly, 30 µl of each serum sample were mixed with 90 µl of binding buffer (PBS) and added to the corresponding well on the equilibrated protein G plate. Following a 45 min incubation the plate flow-through was reapplied to the corresponding position in the purifying plate and incubated again. The flow-through was then saved as “IgG-depleted“ serum. The plate was then washed 4 times with PBS and reconstituted applying 0.1 M glycine ph 2.5. Depletions were verified by IgM and IgG ELISA.

#### Antigen-specific IgG depletion

For the depletion of antigen-specific IgG antibodies, we made use of the high-affinity binding of the double strep-tag encoded in the C-terminal end of the recombinant proteins to MagStrep XT beads (IBA, 2-4090-002). Three depletion rounds were performed to achieve complete depletions. In every round, 20 µl of beads were equilibrated in binding buffer (100 mM Tris-HCl + 150 mM NaCl + 1mM EDTA, pH 8.0) and conjugated with 5 µg of recombinant protein for at least 45 min at 4 °C. Next, unbound protein was removed with a magnetic separator and beads were washed with PBS before adding the serum samples. Usually, IgM depleted-serum samples were diluted in PBS on a final volume of 160 µl (final dilution 1:7.5) and incubated for 1.5 h at RT in a tube rotator. Depletions were verified by IgG ELISA with the respective E protein used for depletion.

#### Antibody Dependent Enhancement of YF17D virus

ADE of YF17D for baseline serum samples was measured using YF17D-Venus infection of THP-1 and K562 cells (cultured in RPMI-1640 10%, 2 mM L-glutamine, and 1 % pen/strep at 37 °C and 5 % CO_2_). In a 96 well plate, seven 1:3 serial dilutions of every test serum sample, starting with a 1:10 dilution, were combined with YF17D-Venus at an MOI of 1 in uncomplemented RPMI-1640. Following 1 h incubation at 37 °C, the serum-virus mix was transferred to a round bottom 96 well plate with 30.000 cells/well. After two hours incubation at 37 °C, cells were washed three times with PBS and then resuspended in R10 medium and left to rest for an additional 62 h. Cells were then stained with a viability dye (Thermo Fischer, L10120), fixed with 4% PFA and acquired in FACS Canto. To account for inter-day and inter-plate variability, every plate carried the same internal control (serum from a TBEV vaccinate donor) and test samples were normalized against it as follows.

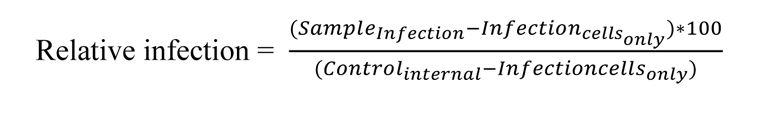

Next, the enhancing titer was quantified as AUC with all serum dilutions as independent variable in GraphPad Prism. Quartile grouping selected top and bottom 25 % as individuals with high and low enhancing titers respectively.

#### Dengue and Zika virus ADE

ADE for dengue and Zika virus was measured for serum samples prior and after YF17D vaccination using previously described DENV and ZIKV reporter virus replicon particles expressing a Gaussia luciferase (45).

In a 96 well plate, five serial dilutions of every test serum sample (1:20, 1:10^2^, 1:10^3^, 1:10^4^, 1:10^5^ for cohort samples or 1:50, 1:500, 1:5.000, 1:50.000 for antigen-specific depleted sera), were combined with VRPs at an MOI of 0.5 in uncomplemented RPMI-1640. Following 1 h incubation at 37 °C, 10.000 K562 cells were added to the serum-virus mix in every well. After two hours incubation at 37 °C, cells were washed three times with PBS, resuspended in R10 medium and then left to rest for an additional 72 h. Supernatants were then collected and stored until measurement. Luciferase activity was quantified in 20 µl of sample using a FLUOstar Omega reader (BMG Labtech). 20 µM of Coelenterazine substrate (Carl Roth, 4094.3) was injected in every well and signal was acquired for 10s as previously described (68).

To account for inter-day and inter-plate variability, every plate carried the same internal control (serum from a TBEV and YF17D vaccinated donor) and test samples were normalized against it as follows.

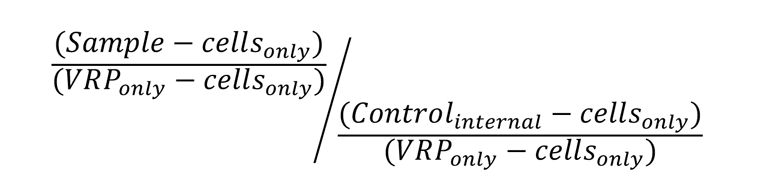

#### B cell EliSpot

B cell EliSpot was used for the quantification of total IgG-secreting, vaccine-specific and epitope-specific plasmablasts and memory B cells present in PBMC samples. Briefly, ethanol-activated polyvinylidene difluoride membrane EliSpot plates (Millipore, MAIPS4510) were coated overnight at 4 °C with 10 µg/ml of capture anti-human IgG (Sigma Aldrich, I2136) in PBS. After being washed with PBS and blocked with R10 for 1 h, cells were added. Cryopreserved PBMC were thawed and allowed to rest for 1 h in warm R10 media (RPMI 1640 supplemented with 10% heat-inactivated FCS, 1% penicillin/streptomycin, and 1% L-glutamine). For the detection of plasmablasts, 100.000 cells/well were added to the plates without further processing. For the memory B cell EliSpot, PBMC were stimulated with 1 µg/ml R848 (SigmaAldrich, SML0196) and 10 ng/ml recombinant human IL-2 (Biolegend, 791904) for 5 days before harvesting and plating of 50-100.000 cells/well. Plates were then incubated at 37 C 5% CO_2_ for 20 h. Uncoated wells were used as negative control and 5.000 cells/well were plated for bulk IgG detection as positive control. Plates were then washed with PBS+0.05% Tween-20, followed by an incubation with 1 µg/ml of recombinant sE WT or dimer-mutants or anti-human IgG-HRP (Jackson ImmunoResearch, 109-035-088, 1:5000) in PBS+0.5% FCS for 2 h at RT. Following thorough washes with PBS+0.05% Tween-20 and PBS 650 mM NaCl, plates were incubated with StrepTactin-HRP (Bio-Rad, 1610380, 1:5000) which binds the Twin Strep-Tag included in the C-terminus of all the recombinant proteins. Plates were finally washed again with PBS+0.05% Tween-20 and spots were developed for 10 minutes with TMB substrate (Mabtech, 3651-10). Plates were rinsed with water and dried before counting the spots with EliSpot Reader ELR04 SR (AIDAutoimmun Diagnostika GmbH, Strassberg, Germany)

## Statistical analyses

Statistical analyses were performed with Prism 8 software (GraphPad) or R and are specified in each corresponding figure caption.

## Acknowledgements

The authors thank Natalie Roeder, Nicole Lichter and Christine Hoerth for technical assistance. The authors also thank Arne Kroidl, Günter Fröschl and Kristina Huber for serving as clinical study investigators. We thank Liz Schultze-Naumburg for support in building up the yellow fever vaccination cohort. We thank Renate Stirner for her assistance with the ELISpot reader. We thank Ritu Mishra and Cell Analysis Core Facility TranslaTUM for their kind help with taking the microscopy pictures for the IIFT assay. We thank Janett Wieseler and Arlen-Celina Lücke for VRP production. We also thank Malena Bestehorn-Willmann and Simon Philipp Stützle for their great support in the implementation of the PRNT assay. We acknowledge the iFlow Core Facility of the University Hospital Munich for assistance with the generation of flow cytometry data. The authors also thank all the cohort participants who voluntarily participated in the study and donated samples. Parts of this work have been performed for the doctoral theses of ASP, FL, SG, EN, MZ, and LL, at the Ludwig-Maximilian University Munich.

## Author Contributions

ASP designed and performed experiments, analyzed and interpreted the data, and wrote the manuscript; FL, SG, EN, EW, MZ and LL performed experiments and analyzed and interpreted the data; FD performed experiments and provided technical assistance; HK provided theoretical assistance and acquired funding; BMK provided critical reagents, acquired funding and helped interpreting the data; GD provided critical reagents and helped interpreting the data; MH and JTS helped to initiate and set up the cohort ; SE supervised the project and acquired funding; ABK acquired funding and supervised the project; MP helped to initiate the cohort, performed vaccinations served as clinical study investigator, acquired funding, and supervised the project; GB-S provided critical reagents, designed experiments, acquired funding and supervised the project; SR conceived the study, set up the cohort, interpreted data, acquired funding, supervised and gave the general direction of the project and wrote the manuscript. All authors critically reviewed the manuscript and approved the final version.

