## Extended Data Figures 1-5 for "Prior flavivirus immunity skews the yellow fever vaccine response to expand cross-reactive antibodies with increased risk of antibody dependent enhancement of Zika and dengue virus infection"

### Extended data Figure 1

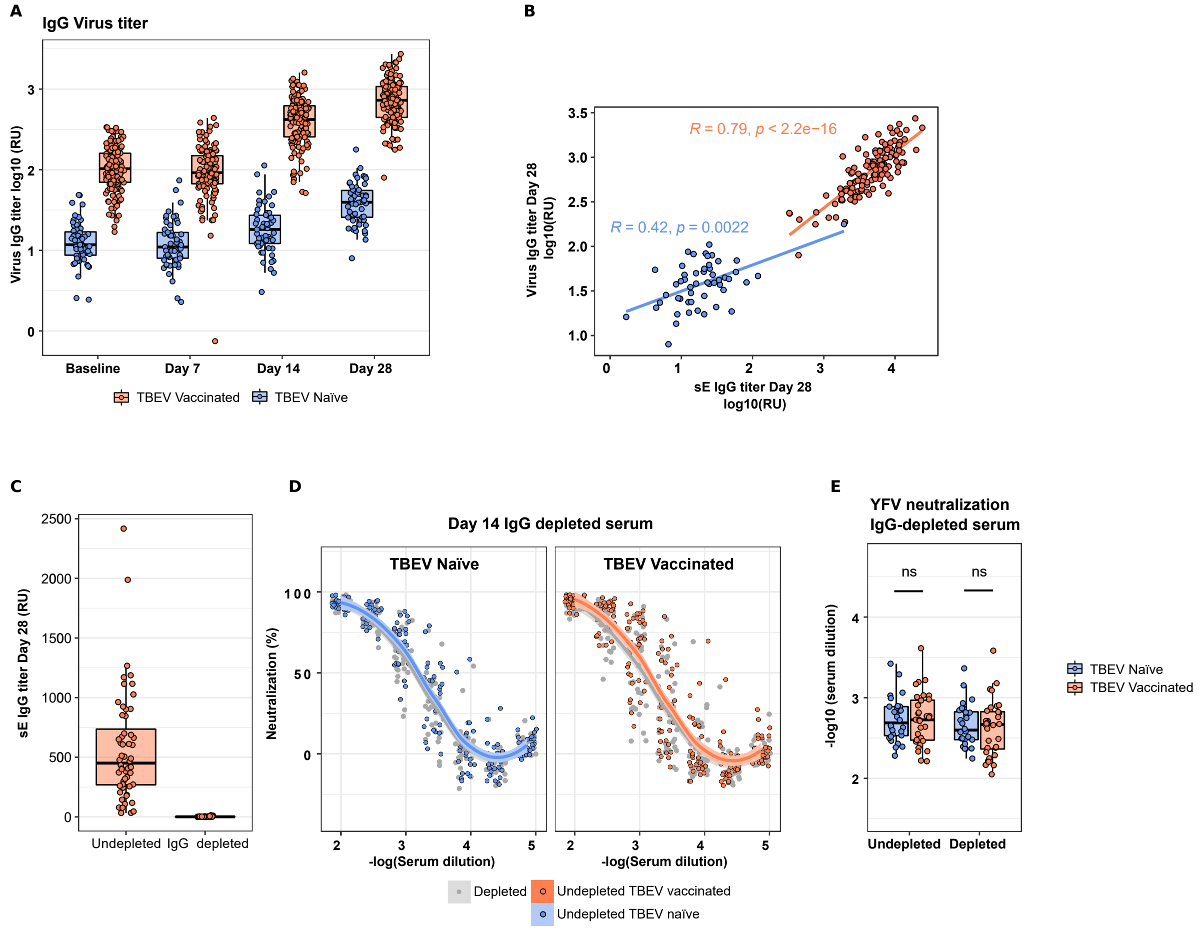

**Extended Data Figure 1**

A) Longitudinal YF17D virion-specific IgG quantification. B) Spearman correlation between the anti-E IgG and anti-YF17D virion titer at day 28. C) IgG depletion validation quantified in an anti-E protein ELISA. D) Neutralization curves of undepleted polyclonal serum and IgG depleted serum (in grey) for TBEV pre-vaccinated (in orange) or naive individuals (in blue). Curve fitting with local regression for n = 45 TBEV vaccinated and n = 19 TBEV naive at day 14 post vaccination E) Quantification of the 80% neutralization cutoff before and after IgG depletions at day 14 post vaccination.

#### Extended Data Figure 2

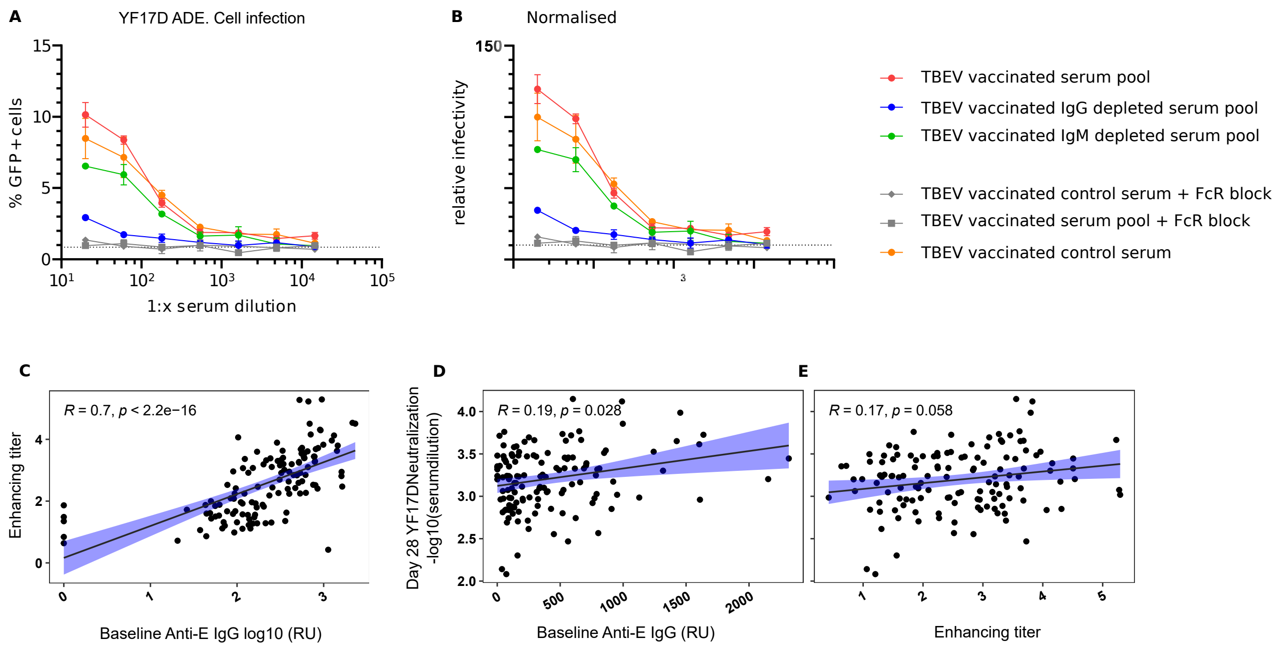

**Extended Data Figure 2**

A-B) YF17D ADE on THP1 cells mediated by undepleted serum or serum with anti-FcγR blocking antibodies and serum depleted for IgG or IgM antibodies. Serum pools of TBEV vaccinated individuals at baseline and the internal control used for normalization across assays were included. Frequency of infected cells (A) was normalised against the internal control as described in methods to obtain the relative infectivity values (B). C-E) Spearmann correlations of the Enhancing titer with anti-E IgGs at baseline and with the IgG and neutralizing titers following YF17D vaccination.

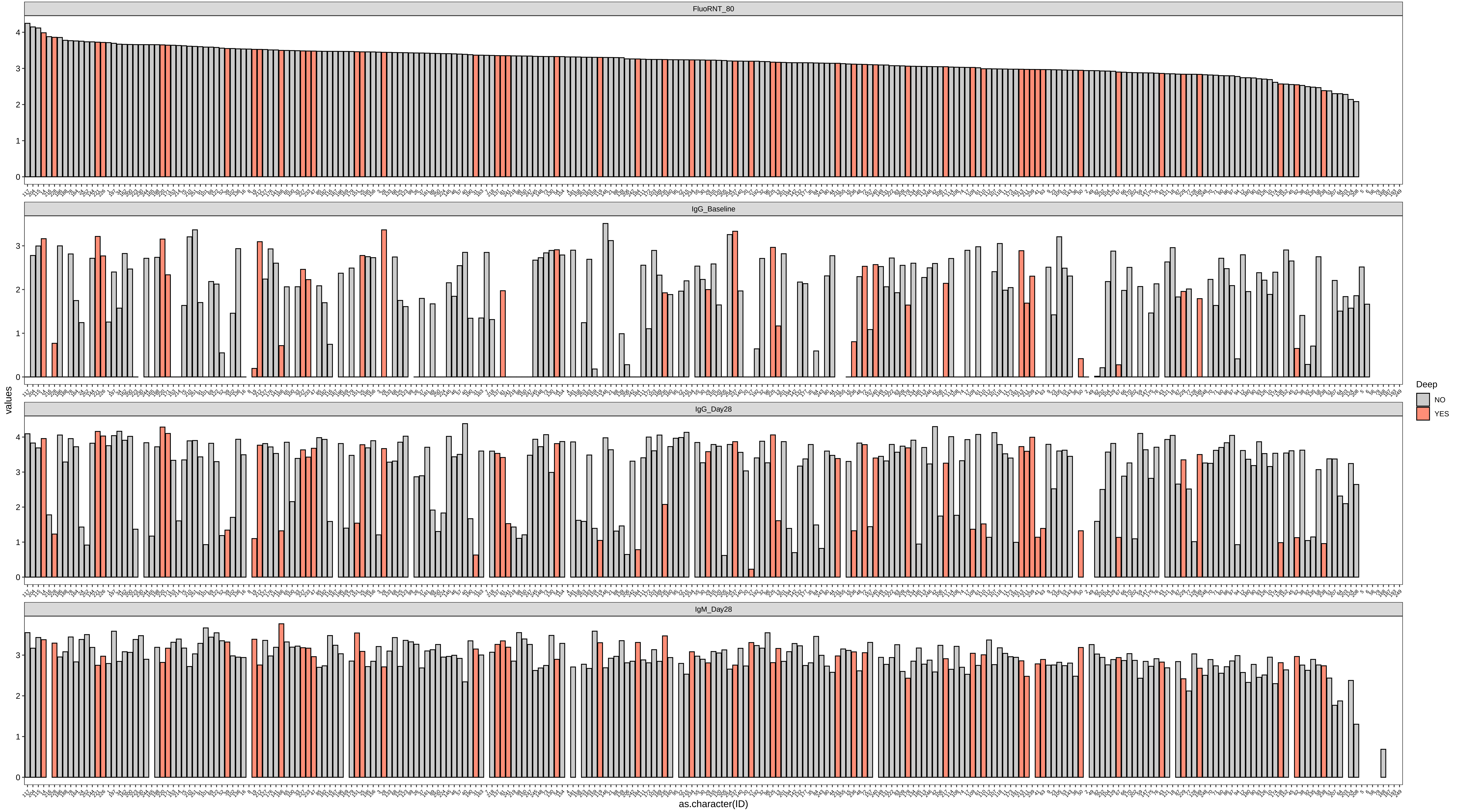
FluoRNT_80
IgG_Baseline
IgG_Day28
IgM_Day28
values
as.character(ID)
Deep
NO
YES

**Extended Data Figure 3**

Individuals selected (in red) for in depth study in cross-reactivity, antigen-specific depletions, DENV and ZIKV ADE, ELISpot and recombinant dimers IgG quantification are representing individuals with different neutralizing capacity and antibody titers after vaccination

#### Extended Data Figure 4

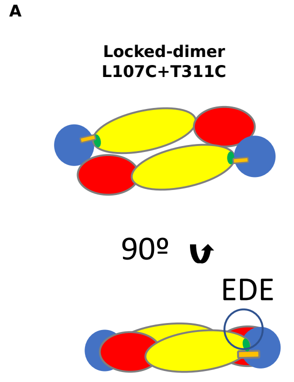


B

| Locked dimer L107C T311C | Expected | Reality |
|---|---|---|
| Epitope specificity | EDE, DI, DII, DIII | DI, DIII |
| Undepleted fraction (remaining specificities) | FLE | EDE, FLE, DII |

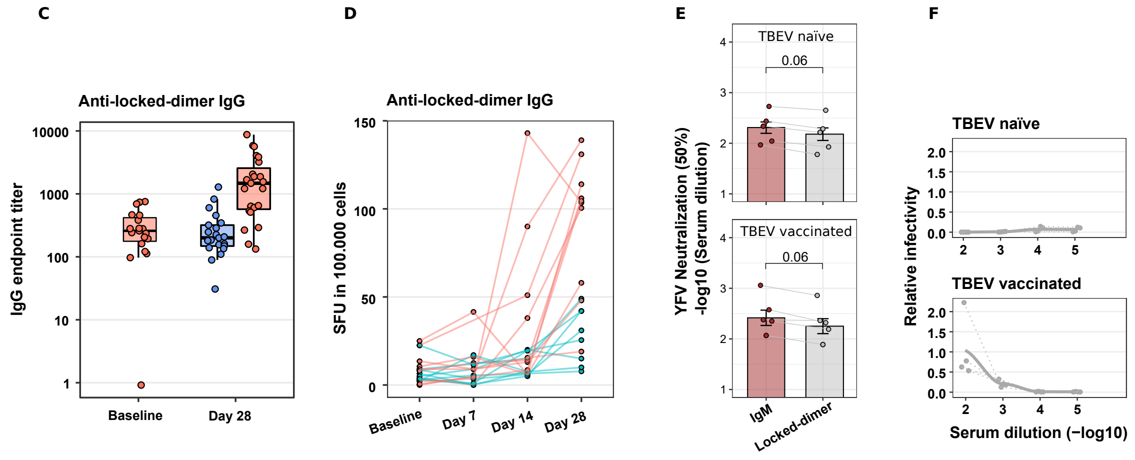

**Extended Data Figure 4.**

A) Representation of the recombinantly produced locked-dimer construct with the disulphide bond formed between the mutated L107C and T311C. B) Table summarizing the expected epitope display for the locked-dimer construct versus the observed real epitope display. Likewise, the table shows the expected remaining specificities after antigen-specific depletions and the actual undepleted fraction. C) Quantification of IgG endpoint titers against locked-dimer protein at baseline and day 28. D) Longitudinal enumeration of IgG-producing B-cells specific for the locked-dimer antigen. Units represent spot-forming units per 100.000 PBMC. E) YF17D neutralization titers (50% cutoff) of IgM depleted and locked-dimer depleted sera. F) Dengue ADE driven by IgM and locked-dimer-specific-IgG depleted sera.

**Extended Data Figure 5**

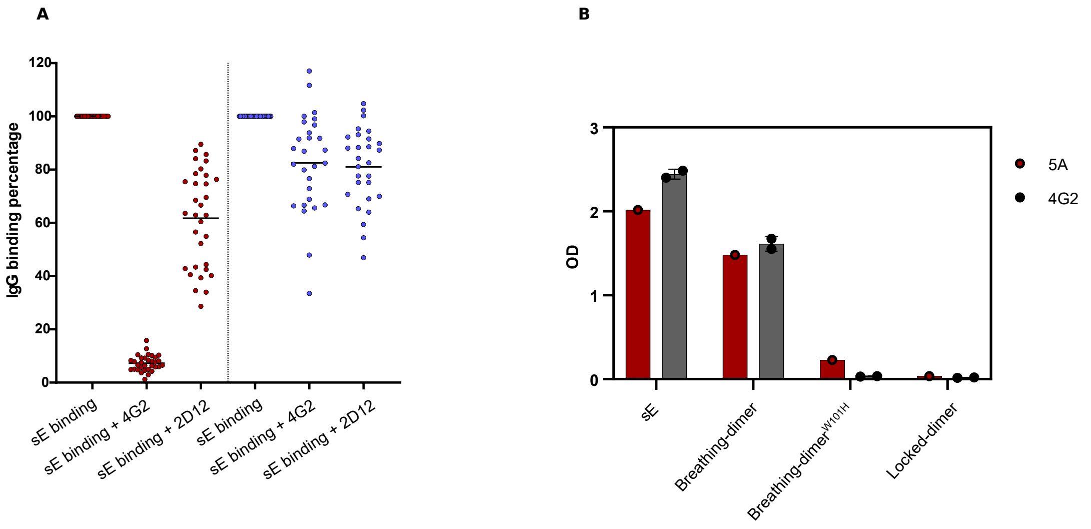

**Extended Data Figure 5**

A) IgG competition binding assay to sE in the presence or absence of competing 2D12 or 4G2 monoclonal antibodies. TBEV pre-vaccinated individuals are depicted in red and TBEV naïve in blue B) ELISA determining binding capacity of the monoclonal 5A and 4G2 antibodies to the plate-coated sE , breathing-dimer, breathing-dimer$^{W101H}$ and locked dimer constructs.
